# Sequence-based coevolutionary prediction of species-specific interactomes

**DOI:** 10.1101/2024.12.27.630483

**Authors:** Miguel Fernandez-Martin, Camila Pontes, Victoria Ruiz-Serra, Alfonso Valencia

**Affiliations:** Barcelona Supercomputing Center, Barcelona, Spain; Universitat Autònoma de Barcelona, Barcelona, Spain; European Commission, Joint Research Centre (JRC), Geel, Belgium; ICREA, Barcelona, Spain

## Abstract

Evolutionarily conserved protein-protein interactions (PPIs) reveal fundamental biological processes. However, predicting protein function solely from these interactions provides an incomplete picture of biological systems. Computational methods for predicting PPIs often struggle due to limited functional annotations in databases, making it difficult to fully understand unique biological systems.. This study introduces ContextMirror2.0 (CM2.0), a coevolution-based method designed to address these limitations. Coevolutionary approaches, unlike supervised machine learning methods, do not rely on labelled datasets, proving valuable in addressing the scarcity of data for species-specific PPIs. While around 40% of CM2.0’s top-1000 predicted PPIs for *E. coli, S. enterica*, and *S. aureus* are present in experimental PPI databases, a comparative analysis of predicted functional communities revealed highly conserved interaction patterns and species-specific interactions. CM2.0 leverages coevolutionary information to explore protein interaction dynamics and understand the functional consequences of species-specific variations. This information can inform further studies on the emergence of novel functions, adaptation to specific environments, and the development of targeted therapies.

## 1 INTRODUCTION

Understanding protein interactions is crucial, yet significant knowledge gaps remain, especially concerning species-specific interactions and those in non-model organisms with poorly annotated proteomes. Computational methods offer valuable alternatives to generate testable hypotheses to cover the knowledge gap on protein complex annotation ^1^.

Current computational methods for predicting complete interactomes are limited. Many machine learning (ML)-based approaches require extensive labelled data, hindering reliable extrapolation beyond a few reference proteomes ^2–4^. Furthermore, the explainability of ML models, particularly deep neural networks, is often limited, and the scarcity of negative examples in experimental PPI databases further complicates training ^5,6^. To address these challenges, we have developed ContextMirror2.0 (CM2.0), a coevolution-based unsupervised method built upon our previous work ^7^. CM2.0 has comparable performance to state-of-the-art coevolution-based methods like Cong et al.’s approach ^8^, while also directly quantifying the contribution of factors such as the number of proteins associated with a given PPI and the strength of the coevolutionary signal, and providing a confidence score for each prediction. This score reflects the reliability of the predicted interaction, offering a measure of certainty based on these contributing factors.

We validated CM2.0 predictions using a combined dataset composed of data from STRING (only high-confidence), IntAct, EcoCyc and Interactome3D. This validation set includes both direct physical interactions and indirect functional interactions (e.g. proteins in the same complex that do not contact directly) and is, therefore, adequate to CM2.0 that, as an evolution-based method, does not differentiate between physical and non-physical associations. However, it is important to highlight that evaluating predicted PPI networks is challenging, even for small bacterial proteomes since a vast number of potential PPIs still lack experimental evidence ^9^.

Our analysis of different bacterial proteomes demonstrates that CM2.0 correctly ranks predicted protein interactions. For a given genome, the top 1000 predictions achieve approximately 40% precision. This accuracy, while moderate, is clearly significant and sufficient for identifying sets of specific PPIs, especially when reinforced by community search of combined multispecies PPI networks as we observed a 3-8% increase in precision for intra-community edges. High-confidence communities are characterised by their presence in multiple species and a high density of precisely predicted pairwise interactions. Interestingly, these communities often involve poorly characterised proteins, offering valuable insights that sometimes constitute the first hypothesis regarding their potential function. Additionally, by comparing interaction networks across species, we identified species-specific interactions worthy of further investigation, including examples, such as proteins from the phn gene community in *E. coli* and a flavoprotein complex in *S. enterica*.

In summary, CM2.0 offers a valuable approach for predicting PPIs, identifying both conserved interaction clusters while also uncovering species-specific interactions with potential biological significance. This information can contribute to the functional annotation of poorly characterised proteins and the identification of new complexes in understudied organisms, as exemplified by the case of *S. aureus* in this study. The associated confidence scores enhance the value of CM2.0 predictions, providing a reliable guide for future experimental and computational research.

## 2 RESULTS

### 2.1 ContextMirror2.0 accurately predicts pairwise PPIs

We applied ContextMirror2.0 (CM2.0), a new high-throughput version of the ContextMirror method ^7^ (see Methods), to predict the interactomes of *E. coli*, *S. enterica*, and *S. aureus*, based on species-specific alignments using as heads the proteins of each one of the three organisms. For each protein pair, CM2.0 calculates an interaction score reflecting coevolutionary signal. This score incorporates the phylogenetic correlation between protein families, corrected for background noise, and the average influence of third proteins’ phylogenetic profiles, that is, proteins that influence the correlation between two given protein interactors (see Methods). Higher scores reflect stronger coevolutionary signals, which is interpreted as an indicator of the quality of the predictions. There is also a positive relationship between phylogenetic correlation and the number of influencing third proteins (**Figure S1**), suggesting that more strongly correlated profiles tend to be influenced by many third proteins. Since this relationship is weaker for low correlations, CM2.0 applies a penalty for protein pairs showing influence from too few third proteins (< 50).

However, the integration of these components into CM2.0’s interaction score could lead to potential biases. Specifically, when a PPI is influenced by only a few third proteins, the interaction score can be inflated by a single strong partial correlation. In contrast, when many third proteins are involved, the interaction score is averaged across all of them, potentially lowering the score but providing a more comprehensive coevolutionary context, especially for PPIs influenced by a small number of third proteins. To reduce the influence of isolated high partial correlations (influences), we introduced a penalization strategy to lower the IS of PPIs influenced by a small number of third proteins (<50). **Figure S1** illustrates the relationship between the correlation of coevolutionary profiles and the number of influencing third proteins. Notably, the curve levels off below 50 third proteins, and statistical tests (See Methods) revealed a significantly higher concentration of false positives (FPs) in this range. Based on these findings, we selected this cutoff for the penalization adjustment.

CM2.0 interaction score (IS) is a number between 0 and 2. The selection of an appropriate IS threshold is crucial when analysing CM2.0 predictions, as it serves as a quality filter. The ideal threshold will depend on the application. A low threshold will include a high number of PPIs of both high- and low-confidence, while a high threshold will include a lower number of high-confidence PPIs. Here, we considered three IS thresholds: 1.6, 1.7, and 1.8, adjusting them as needed to address specific research questions.

**Figure 1A** demonstrates high precision for top-ranked interactions (approximately 40% for the top 1000 predictions), decreasing to around 20% with more predictions across all species. These precision values are reasonable considering the vast number of potential interactions (approximately 9 million for *E. coli)* and potential incompleteness of the ground truth data (precision values for various interaction score thresholds are given in **Table S1**). The precision of our highest-ranked interactions significantly surpasses baseline rates (3.6% for *E. coli*, 2.9% for *S. enterica*, and 2.2% for *S. aureus*), representing a substantial improvement over random chance. This indicates that CM2.0 effectively ranks known interactions while also identifying novel potential interactions.

**Figure 1.**
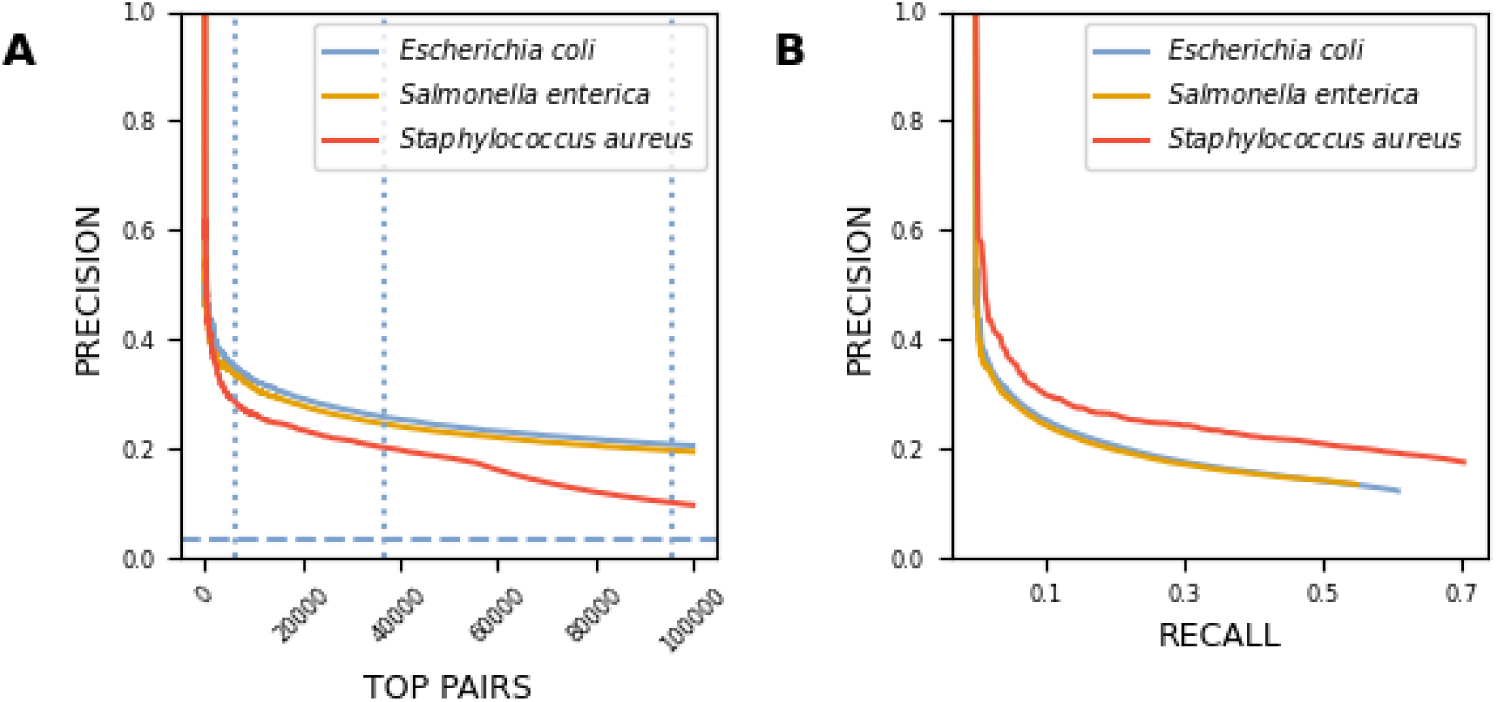
Pairwise accuracy of ContextMirror2.0 predictions. **A)** Precision results for different sets of top-ranked predicted interactions for *E. coli* (blue), *S. enterica* (orange) and *S. aureus* (red). The dashed line corresponds to the precision values that would be expected by chance for *E. coli* (3,6%), *S. enterica* (2.9%) and *S. aureus* (2.2%). The dotted lines indicate the precision corresponding to different interaction scores of 1.8, 1.7 and 1.6 in *E. coli*. B) Precision-recall curve for all species.

Next, we examined the precision-recall trade-off (**Figure 1B**). For *E. coli*, CM2.0 predicted 505,294 PPIs, compared to 312,961 interactions from the STRING database. It is important to note that CM2.0 does not assign scores to all potential PPIs (see Methods). Our results indicate that approximately 20% of the ground truth set was recovered with a precision of roughly 20%, comparable to the performance reported by Cong et al., using residue-level coevolution (DCA) in *E. coli* ^8^, although this approach was computationally more intensive.

We investigated the relationship between interaction scores and paired multiple sequence alignment (MSA) depth, i.e. number of effective sequences. A moderate correlation was observed (0.47 for E. coli, 0.48 for S. enterica, and 0.3 for S. aureus, all with p-values = 0.0), with alignment depth explaining 13% of the variance in interaction scores for *E. coli* and *S. enterica* (16% for *S. aureus*) (**Figure S2**). This indicates that while alignment depth contributes to interaction scores, it is not the only determining factor.

To further characterise some of the PPIs predicted by CM2.0 we modelled their complex structures using AlphaFold (AF) Multimer^10,11^. Our hypothesis was that AF Multimer would produce better structural models for protein pairs with high interaction scores and vice versa, with low-scoring pairs yielding less accurate models. Thus, we modelled a set of 10 high and 10 low interaction score pairs, randomly selected from each organism, resulting in a total of 60 pairs and we evaluated the quality of the modeled complexes using the pDockQ2 score ^12^(see Methods). None of these PPIs are present in the PDB. Three out of 30 tested interactions with high interaction score showed a pDockQ2 score > 0.23, a threshold for acceptable model quality ^12^ (**Figure S3**). Their structural models are represented in **Figure 2**. These results highlight the limitations of AF Multimer in predicting PPIs, particularly when the interactions are not of a physical type. In such cases, additional aspects need to be considered, as covered by CM2.0. However, AF Multimer could still be useful for making distinctions among interactions, given that 3 out of 17 true positive interactions tested did surpass the pDockQ2 threshold.

**Figure 2.**
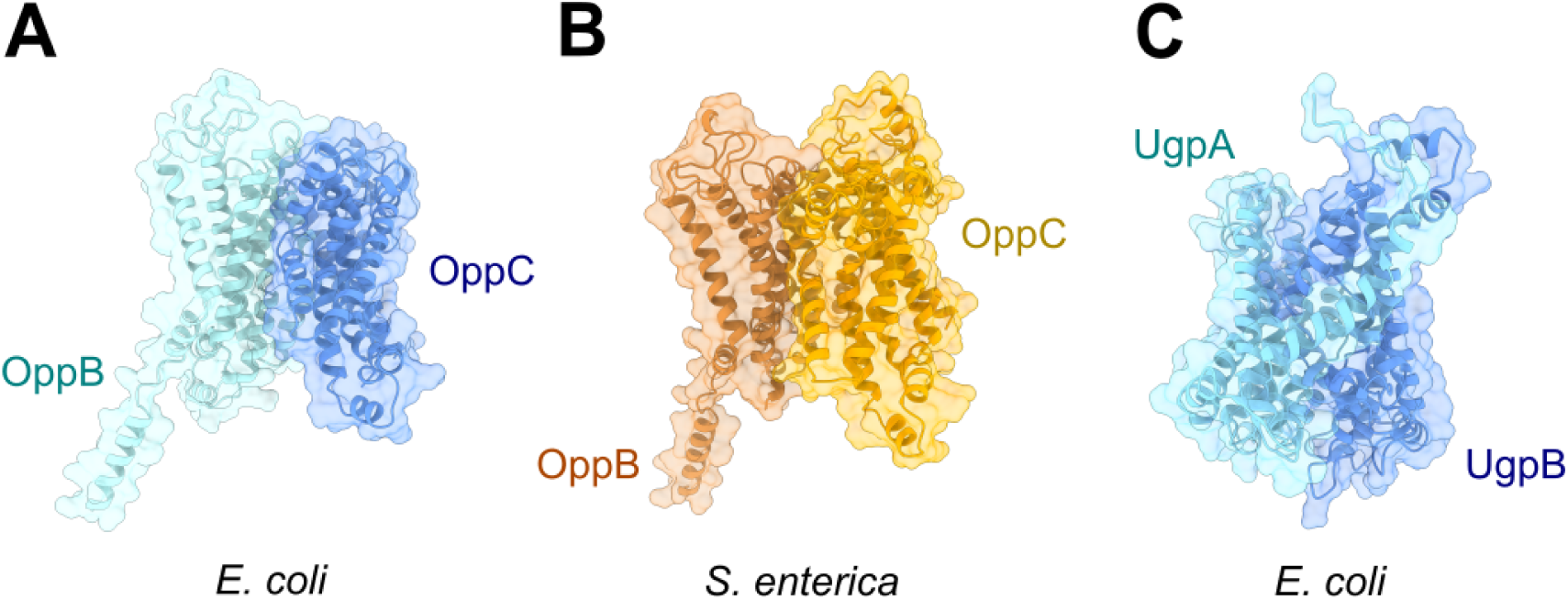
AlphaFold Multimer models with highest min pDockQ2. **A)** OppB (P0AFH2) - OppC (P0AFH6) Oligopeptide transport system in *E coli*. B) OppB (P08005) - OppC (P08006), Oligopeptide transport system permease protein OppB-OppC in *S. enterica*. C) UgpA (P10905) - UgpE (P10906) glycerol uptake system in *E. coli.* None of the produced models are in PDB.

Collectively, the results presented in this section show that CM2.0 tends to consistently assign high scores to true positive PPIs across different species, with an accuracy over ten times higher than what would be expected by chance and comparable to other similar methods. In the following sections, we will investigate how the pairwise PPIs predicted for individual organisms can be leveraged to identify common and species-specific protein functional communities.

### 2.2 Species-specific functional communities can be identified within a multispecies PPI network

In this section, we evaluated CM2.0’s predictive accuracy in the context of conserved and species-specific protein-protein interactions (PPIs). To achieve this, we identified common and distinct PPIs by mapping homologous sequences across species (see Methods).

As expected, the Gram-negative bacteria *E. coli* and *S. enterica* shared a high proportion of pairwise interactions (70.0%), whereas the more distantly related Gram-positive *S. aureus* shared fewer interactions with both (40.7% and 48.3%, respectively) (**Figure 3**), a finding consistent with their phylogenetic distance and functional differences ^13^. High-confidence predictions (i.e., higher interaction scores) showed greater conservation across species (**Figure S4**), suggesting that conserved PPIs tend to involve ubiquitous bacterial proteins. Conversely, species-specific PPIs, being less well-characterised, rely on less diverse alignments, potentially leading to weaker coevolutionary signals. This reflects a precision-recall trade-off when predicting species-specific interactions.

**Figure 3.**
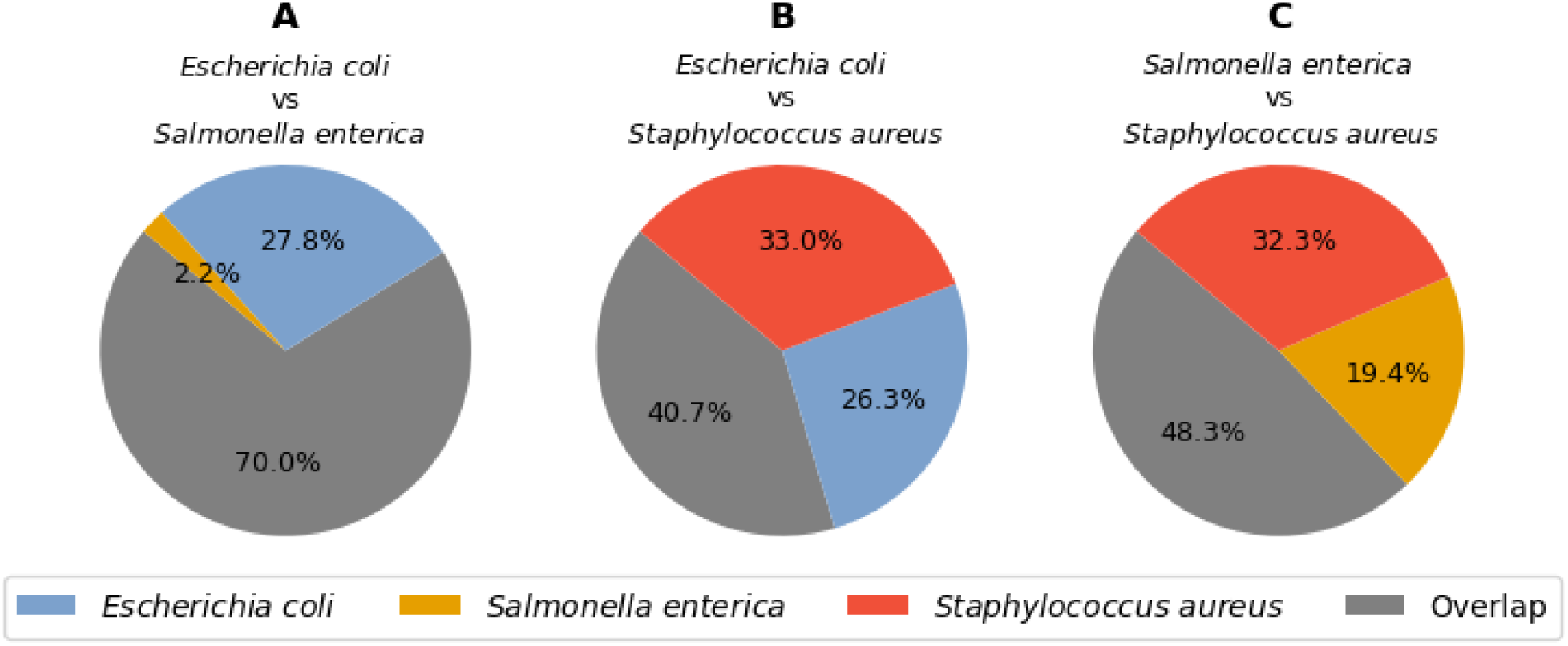
Percentage of shared PPIs between species. Panels A, B and C show the PPI overlap between *E. coli* and *S. enterica* (n = 467505), *E. coli* and *S. aureus* (n = 40403), and *S. enterica* and *S. aureus* (n = 53580), respectively. Shared interactions between species are shown in gray, *E. coli* unique interactions are shown in blue, *S. enterica* unique interactions are shown in orange, and *S. aureus* unique interactions are shown in red.

Next, we constructed PPI networks for the two better-characterised species, *E. coli* and *S. enterica,* with edges weighted according to their interaction score (IS). We then combined these two networks by matching homologous protein nodes. This was done considering different IS thresholds, i.e. IS > 1.8, IS > 1.7 and IS > 1.6. For the strictest threshold (IS > 1.8) for both species, precision was around 35%, and the combined network contained 531 nodes and 1492 edges (see **Table S2** for thresholds 1.6 and 1.7).

To identify functional associations between groups of proteins, we predicted communities in the combined network using the Louvain algorithm with different initial random states (see Methods). **Figure S5** shows the communities obtained on the run with highest overall intra-community precision, while the results presented in the following are averages from multiple runs. **Figure 4** shows the average percentages of true positives (TPs) and false positives (FPs) within communities for the combined networks computed with different IS thresholds. The results show that as the IS threshold increases, the percentage of TPs rises and FPs decreases in both species, as expected. The proportion of predicted interactions that are TPs in one species and FPs in another remains low and constant across IS thresholds.

**Figure 4.**
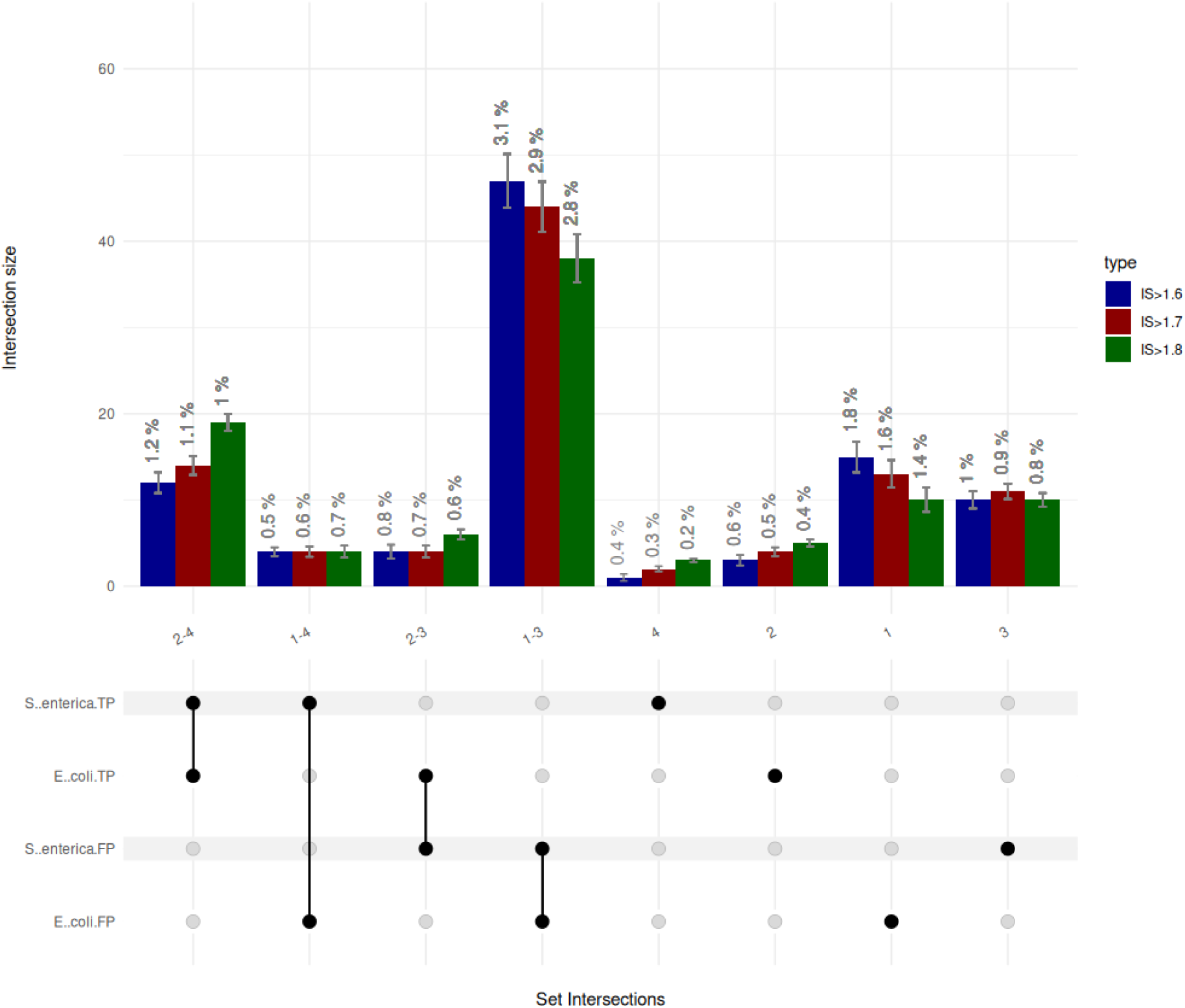
Average number of intra-community edges of different types in the combined PPI network of *E. coli* and *S*. *enterica*, as inferred by CM2.0. The first four sets (2-4, 1-4, 2-3 and 1-3) represent common PPIs while the last four sets (4, 2, 1 and 3) represent species-specific PPIs. The predicted PPIs are separated according to their validation status in the groundtruth PPI sets as a known interaction or true positive (TP), or as an unknown interaction or false positive (FP). The average number of intra-communities edges belonging to each set is shown for different interaction score (IS) thresholds: IS > 1.6, IS > 1.7 and IS > 1.8. Communities were determined using the Louvain algorithm with different initial random states.

This analysis identified three sets of interesting PPIs: (1) PPIs reported in one species but not in the other (sets 1-4 and 2-3 in **Figure 4),** (2) unreported PPIs identified in both species (set 1-3 in **Figure 4**), and (3) both reported and unreported species-specific PPIs (sets 4, 2, 1 and 3 in **Figure 4**). Specifically, we identified around 104 potentially novel PPIs in *S. enterica* (reported in *E. coli*), 78 novel PPIs in *E. coli* (confirmed in *S. enterica*), 610 previously uncharacterized PPIs shared by both species, and 249 and 226 potentially novel species-specific PPIs in *E. coli* and *S. enterica*, respectively. Similar results, though with lower precision, were observed at lower interaction score thresholds (**Table S2, Figures S6 and S7**).

Finally, the results show that intra-community precision slightly exceeded pairwise precision on average at all thresholds (e.g., 36.9% vs. 34.8% for *E. coli*, IS > 1.8), an indication that community-based analysis constitutes a reliable approach for predicting protein functional communities. When comparing species-specific vs. common PPIs, intra-community precision was lower for species-specific PPIs: 34,6% *vs.* 37.4% for *E. coli* and 27% *vs.* 35% for *S. enterica*. This reinforces the conclusion that, perhaps not surprisingly, species-specific PPIs identified by CM2.0 tend to correspond to less well-characterized interactions than those common across species.

Collectively, the results presented in this section show how pairwise interaction scores generated by CM2.0 can be leveraged to identify shared functional communities and potential species-specific relations. In the next section, we provide further details on some of the species-specific functional communities identified in *E. coli* and *S. enterica*.

### 2.3 Examples of species-specific functional communities

To further explore examples of functional protein communities with species-specific PPIs, we highlight two communities that are entirely species-specific. The first is the *phn* gene community, which drives the conversion of methylphosphonate into phosphate and methane ^14^, was partially identified in *E. coli,* but not in *S. enterica* (**Figure 5A**). This is because the complex structure in *S. enterica* comprises different proteins ^15^ and the coevolutionary signals between them were not captured by our current set of parameters. The second community involves three electron transfer flavoproteins exclusive to *S. enterica* (STM0855, STM0856 and STM0858) ^16^ (**Figure 5B**). This *S. enterica*-specific community also includes yibF (STM3684), a glutathione S-transferase, and rfbI (STM2093), involved in O antigen modification as a putative ortholog of gtrB ^17^. The identification of these communities highlights the capacity of PPI predictions to uncover complexes shaped by evolutionary pressures and ecological contexts specific to, in this case, *S. enterica*. RfbI’s role in modifying the O antigen could be particularly relevant to host-pathogen interactions. Similarly, the interaction with yibF suggests a coordinated response to oxidative stress or other metabolic challenges unique to *S. enterica*.

**Figure 5.**
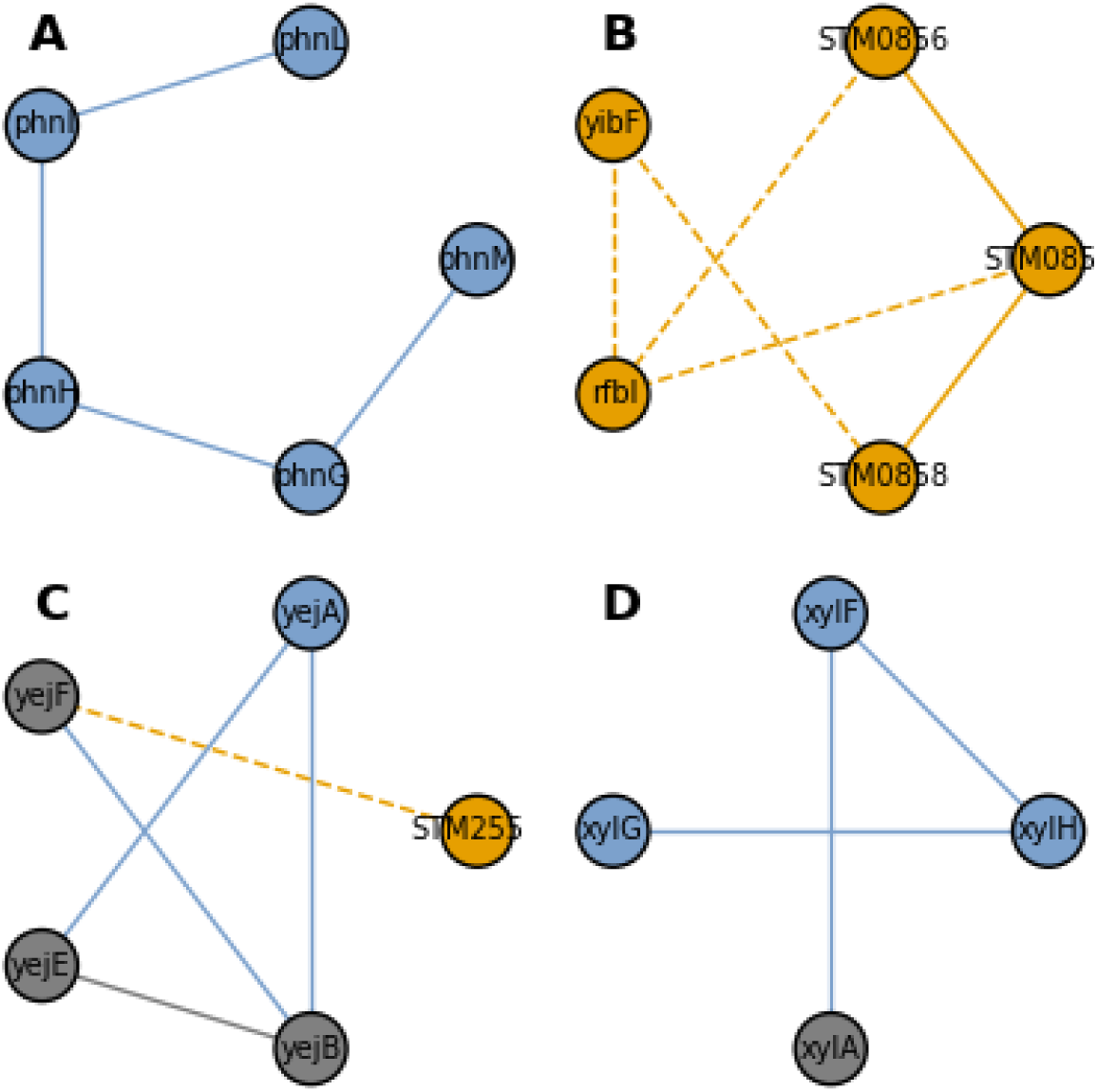
Examples of protein communities detected in the combined PPI network of *E. coli* and *S. enterica* at the threshold of IS > 1.7 containing species-specific PPIs. **A**) A community containing proteins from the *phn* gene cluster. **B)** A community containing three electron transfer flavoproteins (STM0855, STM0856 and STM0858). **C)** A community containing proteins encoded by the *yej* gene cluster. **D)** A community containing proteins encoded by genes form the *xyl* operon. The gray nodes/edges are common to both species, while the blue nodes/edges are only present in *E. coli* and the orange nodes/edges are only present in *S. enterica*.

Besides fully species-specific communities, there are species-specific PPIs within conserved functional communities. One example is a transporter complex composed of yejA, yejB, yejE and yejF (**Figure 5C**), which specifically recognizes the peptide component of Microcin C ^18^, an antibiotic produced by various microorganisms. These genes, present in both *E. coli* and *S. enterica,* confer resistance to antimicrobial peptides and contribute to virulence ^19^. In the identified community, all proteins from the yejABEF complex are present. Notably, we observed a species-specific interaction involving an additional node exclusive to *S. enterica* (STM2551), which forms an uncharacterized interaction with YejF. This interaction suggests functional divergence between the transport systems in *S. enterica* and *E. coli*, as both the Yej complex and STM2551’s closest homolog (E-value = 5.8e-9) in *E. coli* (P76425 - rcnA), a nickel-cobalt efflux pump ^20^, are associated with the ATP-binding cassette (ABC) transporter family, involved in oligopeptide transport (e.g. microcin C) and nickel/cobalt homeostasis (KEGG PATHWAY: ko02010).

Finally, **Figure 5D** shows a community encompassing interactions between proteins encoded by the genes of the *xyl* operon, a key genetic component found in various bacteria that facilitates the metabolism of D-xylose ^21^. While both *E. coli* and *S. enterica* have the xylA gene, *S. enterica* lacks the xylFGH transporter genes ^22^, suggesting an alternative xylose transportation and metabolism mechanisms in this species. The presence of xylA in both species and its capability to operate independently highlight its significance in bacterial metabolism ^21^.

Highly conserved protein complexes, such as the flagellar assembly (*flg*), the ABC-transporter cassette (*ugp*), and the NADH:ubiquinone oxidoreductase (*nuo*) complexes (**Figure S8**), were accurately identified across various interaction score thresholds. Their near-complete conservation of edges between *E. coli* and *S. enterica* reflects their essential roles in Gram-negative bacterial function. These examples show how CM2.0’s can effectively filter predicted networks to identify biologically meaningful interactions. This approach offers valuable insights into species-specific adaptations and the intricate interplay between bacterial genomics and metabolism.

2.4 Multispecies PPI networks can be used to infer the function of unannotated proteins

To conclude our analyses, we selected well-characterized functional communities in *E. coli* and *S. enterica* (e.g., mur, pur, accb/fab) to showcase how CM2.0 predictions can enhance functional annotation of proteins in less-annotated species like *S. aureus.* We retrieved a list of interactors from EcoCyc ^23^ to reveal the predicted topology of these communities. This resulted in subnetworks of proteins encoded by well-annotated proteins in *E. coli* and *S. enterica,* which we compared with predictions in *S. aureus*. To refine annotations and discover additional functionally related proteins, we used a list of interactors as a proxy for identifying functional communities, moving beyond traditional operon-based approaches.

In **Figure 6A**, the conserved *mur* operon (murABCDEFI)^24^ is shown. As expected, interactions between these proteins were identified across all species at relaxed thresholds and only partially at stricter thresholds (**Figure S9**), indicating a very well-conserved coevolutionary signal. murG, the last enzyme in the peptidoglycan synthesis pathway ^25^, is crucial for bacterial cell wall biosynthesis. Unlike predictions for *E. coli* and *S. enterica*, murG is absent in the predicted interactome of *S. aureus*. However, the node WP_000160904.1 forms significant interactions with murA and murF, highlighting its central role in bacterial cell wall synthesis. In fact, there is a record of this sequence (WP_000160904.1) identified as a multispecies murG for all *Staphylococcus* species. Since murG is a promising antibiotic target ^26^, understanding differences in interaction patterns across these subnetworks could reveal potential vulnerabilities in the mur enzyme complex and provide valuable insights for developing antibacterial strategies.

**Figure 6.**
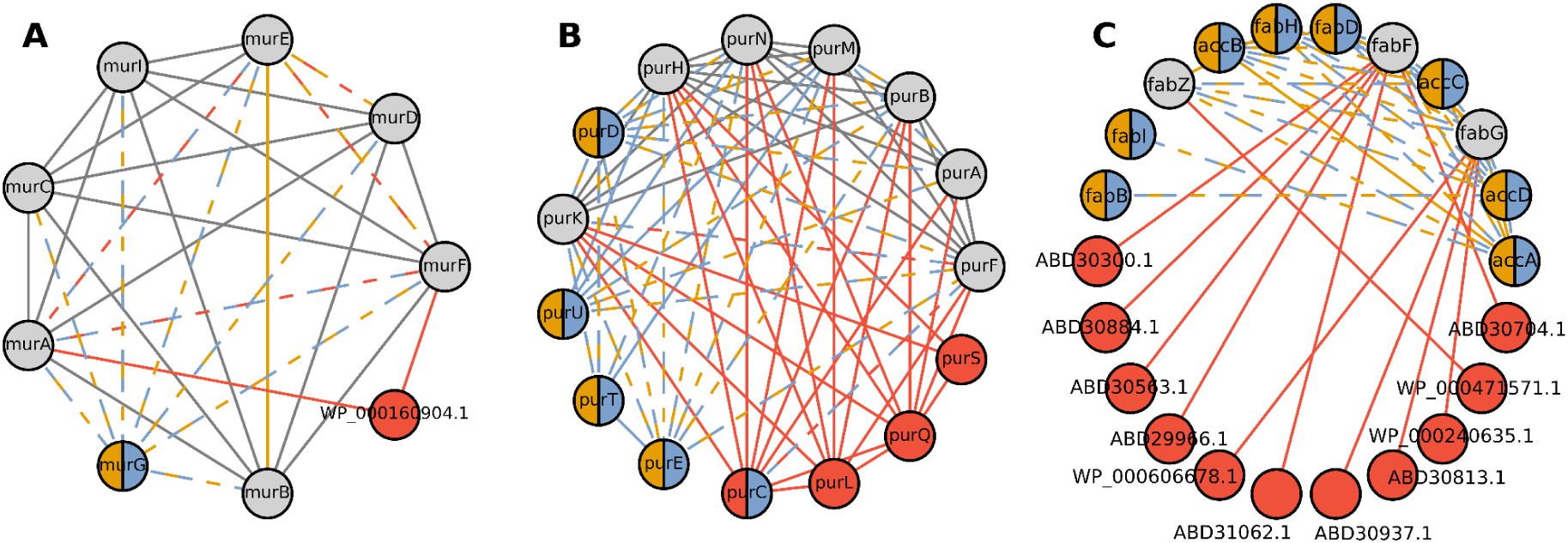
Predicted protein-protein interactions (PPIs) for the mur, pur, and acc+fab complexes in *E. coli*, *S. enterica*, and *S. aureus*. Interactions were filtered using the following interaction thresholds: **A)** 1.2 for the mur complex, **B)** 1.2 for the pur complex, and **C)** 1.7 for the acc+fab complex. Shared nodes and edges are shown in gray, *E. coli* predictions in blue, *S. enterica* predictions in orange, and *S. aureus* predictions in red.

Focusing on the *pur* family ^27,28^, involved in the purine biosynthesis, we identified a highly conserved cluster of proteins (purABCFHKMN) shared among the three species. This community was fully detected at a relaxed threshold of 1.2 (**Figure 6B**) but only partially at stricter thresholds (**Figure S10**). **Figure 6B** shows that some nodes and edges (purDETU) are shared among Gram-negative bacteria, while purL, purQ, and purS are unique to the predicted interactome of *S. aureus*. These proteins constitute the phosphoribosylformylglycinamidine synthase complex (FGAR-AT complex / FGAM synthase complex). In Gram-positive bacteria and archaea, the FGAR-AT complex has three unique components (purS, purQ, purL), absent in Gram-negative bacteria^29^. The CM2.0 approach successfully identified these proteins only in the predicted network of *S. aureus*. Since the purine biosynthesis pathway is a promising antimicrobial target ^30^, this analysis could enhance the development of antibiotic treatments, particularly for Gram-positive bacteria with limited annotation.

As a final example, we examined interactions between proteins from the *fab* and the *acc* operons ^31,32^ (**Figure S11**). **Figure 6C** shows that PPIs involving accABCD proteins were only retrieved for *E. coli* and *S. enterica*. However, these proteins are predicted to interact with nodes in the fab complex, also predicted in the poorly annotated proteome of *S. aureus*. This “bridge” node revealed a novel interaction in S. aureus: WP_000471571.1, potentially linked to the acc complex through its shared interaction with fabZ. Homology searches revealed that WP_000471571.1 matches the accD sequence (Q2YTE0) in *S. aureus* strain bovine RF122 / ET3-1, reported in the STRING database to interact with fabZ with a confidence score of 0.982 for *S. aureus* strain NCTC 8325 / PS 47.

We also identified several proteins of unknown function interacting with fabF. For example, ABD30300.1 likely encodes fatty acid kinase subunit A (fakA), which regulates virulence factor transcription and fatty acid incorporation into phospholipids ^33,34^. Similarly, ABD30884.1 likely encodes pepP, a Xaa-Pro aminopeptidase and known virulence factor in *Campylobacter jejuni* ^35^. ABD30563.1 may correspond to ypjA, an adhesin involved in adhesion and biofilm formation in *E. coli* ^36^, making it a promising candidate for *S. aureus* research. Another interesting interaction involves fabF and ABD29966.1, likely rnmV, a conserved 5S ribosomal maturase in low G+C Gram-positive bacteria ^37,38^. Other notable interactions include ABD31062.1, a protein predicted to be membrane-associated in *E. coli* ^39^, and ABD30304.1, identified as a malonyl CoA-acyl carrier protein transacylase corresponding to fabD. These findings suggest diverse biological roles for fabF in *S. aureus* and merit further investigation.

Finally, fabG interacts with four unmapped proteins in *S. aureus.*, namely WP_000606678.1, identified as Tripeptidase T (pepT) and reported for virulence importance ^40^; ABD30937.1 (contains oxidoreductase HI0933 domain); WP_000240635.1, identified as a chaperonin whose inhibition is a promising antibiotic treatment ^41^; and ABD30813.1, very likely spoT, a protein that increases survival under stress conditions in *E. coli* ^42^.

This section illustrates how multispecies PPIs inferred based on pairwise interactions predicted by CM2.0 can be used to infer the functional role of uncharacterized or unannotated proteins, which is particularly useful for poorly characterized organisms.

## 3 DISCUSSION

Understanding the composition of protein complexes is crucial, yet there remains a significant knowledge gap in this area, particularly with respect to poorly annotated proteomes ^1^. Current computational approaches, including homology-based annotation ^43^ and ML algorithms, face considerable limitations when dealing with poorly annotated proteomes, hindering the prediction of new functions and the identification of species-specific interactions ^2–4^. Moreover, ML approaches sometimes lack interpretability, and might suffer from the lack of negative examples for training^6^. ContextMirror2.0 (CM2.0) offers a suitable alternative that does not require supervision and can produce functional annotations at the single-species level based on MSAs. As a redesign and new implementation of our previous work ^7^, the CM2.0 algorithm shows comparable performance to other coevolution-based approaches like Cong et al.’s method ^8^ and additionally provides information on the significance of the predicted complexes.

The limitation of the evolution based methods is their intrinsic incapacity to differentiate interactions by their nature, and cannot separate functional and physical interactions. Indeed, we did not detect a significant correlation with the potential formation of physical complexes, as assessed by the quality of the corresponding structural models (**Figure S3**). For instance, the AF structure predicted for a CM2.0 high-scoring *S. aureus* PPI, and interaction validated in experimental databases, had a very low pDockQ2. Inversely, the structure predicted for one of the *E. coli* PPIs with low CM2.0 IS, a potential True Negative since it is not confirmed in the experimental databases, has a very high pdockQ2 score (**Figure S12**), a result that is very difficult to interpret. In any case, a full validation of the predicted complexes against structural predictions will remain a challenge dependent on the evolution of current methods ^10,11^, as pointed out in the recent CASP meeting ^44,45^. To validate CM2.0, we used interaction annotations from different PPI databases alongside experimental data, treating functional and physical interactions equivalently.

When analyzing bacterial proteomes, CM2.0 effectively prioritized true protein interactions, with around 40% of the top 1000 predictions matching known annotations from databases. These pairwise PPI predictions were leveraged to build multispecies PPI networks and automatically identify potential protein communities based on the pairwise interactions. Most of the high-confidence communities are present in multiple species and have a high density of interactions confirmed in databases. Some of these involve poorly annotated proteins, offering a first clue about their potential functions. Interestingly, we found a number of communities that are entirely species-specific. For example, the identification of the *phn* gene community in *E. coli* and the electron transfer flavoproteins in *S. enterica* illustrate how CM2.0 can uncover complexes that are relevant to specific metabolic and defensive mechanisms, pointing to host-pathogen interactions and unique metabolic challenges associated to different species. Unlike traditional approaches that rely on operon information to infer functional relationships, we focused on identifying functional communities through protein interactors, using them as a dynamic proxy for relatedness. This shift allows for a more flexible understanding of cellular organization, enabling the discovery of conserved functional relationships across species without the constraints of static gene clusters ^46^. By emphasizing functional communities rather than fixed operon structures, our approach enhances the ability to uncover novel and context-specific interactions, providing a more dynamic and comprehensive view of cellular functions.

To summarize, CM2.0 provides an effective tool for predicting protein interactions, uncovering both conserved clusters that in some cases include proteins that were poorly annotated until now and, on the other extreme, predict potentially interesting species-specific interactions and possible protein communities. CM2.0 can contribute to the annotation of orphan proteins and the identification of new complexes in understudied organisms. Our predictions have an associated score that correlates with the reliability of the interactions and can serve to guide future experimental and computational research. CM2.0 opens the door to future applications extending the range of predictions to less well-known species, following the example of *S. aureus* used in the paper.

## 4 METHODS

### 4.1 ContextMirror2.0

The ContextMirror method (**Figure 7**) comprises 3 steps. First, an initial coevolutionary network is generated by calculating the tree similarities for every protein pair in the input file. Next, an optimized coevolution measure is derived for each protein pair by analyzing their coevolutionary patterns relative to all other proteins. Finally, the influence of additional proteins on the coevolution of a given pair is assessed. Each step is described in detail below.

**Figure 7.**
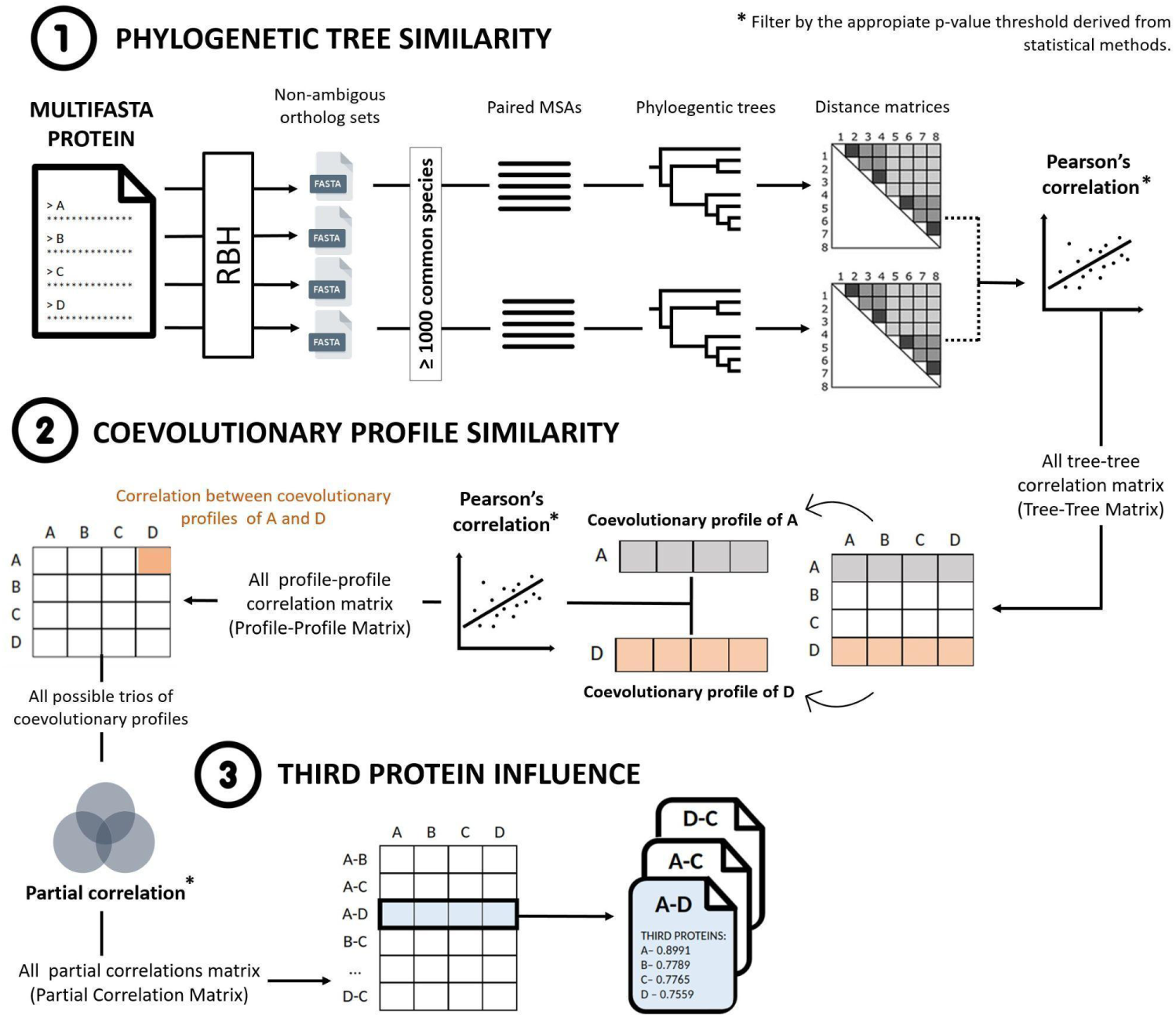
ContextMirror2.0 pipeline. The figure illustrates the overall workflow of the ContextMirror2.0 pipeline, with key steps outlined. Detailed descriptions of steps 1, 2, and 3 are provided in the Methods section (section 4.1, steps 1, 2, and 3, respectively).

#### Step1: All vs. all tree similarities (Tree-tree correlation)

The starting point of ContextMirror is the computation of a network of pairwise tree similarities, that is, the similarity between all possible pairs of phylogenetic trees derived from a given set of proteins (e.g. *E. coli’*s proteome). To obtain these tree similarities, we use the MirrorTree approach ^47^. For each protein, an ortholog set is retrieved using the standard Blast Reciprocal Best Hit (RBH) approach ^48^ with a cutoff e-value of 10^-5^, requiring an alignment of at least 70% of the query protein length, with a minimum 60% query coverage and at least a 30% of sequence identity. The forward Blast step is performed on the full NCBI’s non-redundant database (25/03/2022), while the reverse Blast is performed against the original input set of sequences.

Protein pairs are then filtered to contain a minimum number of coinciding species of their ortholog sets, which in our benchmarking, is a minimum of 1000 common species (**Table S3**). In the following step, paired multiple sequence alignments (MSAs), i.e. MSAs containing only sequences from the coinciding species between the two ortholog sets are produced. Finally, a distance matrix is obtained from every MSA using the McLachlan amino acid residue homology matrix ^49^ and the similarity between the two protein families is calculated as the Pearson’s linear correlation between the two matrices ^50^ as follows:

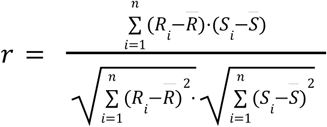

where *n* is the number of elements of the matrices, *n* = (*N*^2^ − *N*)/2, N is the number of sequences in the MSAs, R_i_ and S_i_ are the pairwise distances between proteins in the first and second MSAs, respectively, and 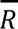 and 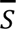 are the means of *R_i_* and *S_i_*, respectively.

Each correlation value *r* is considered significant based on a p-value cut-off of 10^-5^. As a result, a square matrix containing all tree-tree significant correlation values (raw tree similarities) is obtained.

#### Step2: All vs all coevolutionary profile similarities (Profile-profile correlation)

If protein pairs are treated as independent interactors, a high rate of false positives is expected ^51^. To avoid that, the full matrix of coevolutionary information is used to evaluate the coherence of the coevolutionary signal between a pair of protein families. In other words, protein pairs will only be considered as interacting if the comparison of the coevolutionary contexts are similar, i.e. if their pattern of coevolution with other proteins is similar. To evaluate that, we calculate the Pearson correlation between the coevolutionary profiles of the trees. A coevolutionary profile is a vector containing the correlation between a protein and all other proteins in the input set, i.e., a specific row/column of the matrix that was generated in the first step (MirrorTree). The updated correlation coefficient *r’* is then calculated as follows:

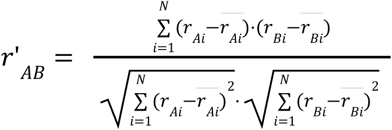

where N is the number of proteins with significant raw tree correlation (only proteins that are significantly correlated to both potential interactors in the pair are considered), r_Ai_ and r_Bi_ are elements of the coevolutionary profiles of A and B, respectively, and 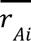 and 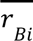 are the means of r_Ai_ and r_Bi_ Since this is done for all possible pairs of proteins, the result of this step is a NxN matrix where the correlation between all pairs of significantly correlated coevolutionary profiles is included.

In this step, given the high number of simultaneous statistical tests being made, a phenomenon known as the multiple testing effect can occur, where an event with a low probability might appear significant due to the sheer number of inferences ^52^. To correct for this effect, a Bonferroni correction is applied, which is a statistical adjustment that reduces the chance of obtaining false-positive results by lowering the threshold for statistical significance in proportion to the number of comparisons. After the Bonferroni correction the p-value threshold in this matrix is set to 10^-11^, making the criteria for significance more stringent and thus reducing the likelihood of false positives.

#### Step3: Influence of all profiles in every profile-profile correlation (Partial correlation)

In this step, it is analysed if the coevolutionary signal between two proteins could be attributed to broad evolutionary tendencies involving many proteins. If correlation is not attributable to any other third protein, it means the signal is highly specific, and thus a probable indicator of a close functional and structural relationship. However, the components of protein complexes influence one another and therefore, patterns of third-protein influence will reveal relationships within these complexes, and with others as well.

To separate specific from general coevolution, the partial correlation coefficient between every pair of proteins and the rest of the proteome is calculated. The partial correlation coefficient between two proteins (A and B) given a third protein (N) is calculated as follows:

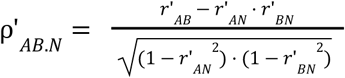

where ρ’_AB.N_ is the partial correlation between the coevolutionary profiles of proteins A and B with a constant coevolutionary profile of protein N, and r’_AB_, r’_AN_, and r’_BN_ are the Pearson’s correlations between the coevolutionary profiles of proteins A and B, A and N, and B and N, respectively. In this step, only partial correlation with a p-value smaller or equal to 10^-6^ were considered significant.

As a result, for each protein pair, a list of partial correlations with third proteins is produced. This list displays the number of third proteins that influence the correlation between the coevolutionary profiles of a pair of proteins, and the magnitude of this influence, represented by the partial correlation coefficient. This way, a protein pair that is influenced by a few or none third proteins or with low partial correlations could be considered as a highly specific correlation. By setting a correlation cut-off, we can extract the number of third proteins influencing the interaction of a protein pair and identify groups of proteins influenced by the same third profiles, indicative of functional relatedness.

The final product of this implementation is a ranked list of predicted protein-protein interactions (PPIs). These PPIs are ranked by the level of confidence that they are true interactions, rather than artifacts produced by the implementation process. This ranking system is based on the integration of coevolutionary information obtained from each of the steps. During the first step, we reduce all possible combinations of proteins to those that share orthologs in at least a 1000 species and show a significant correlation between their evolutionary histories. The second step produces a correlation value for each of the considered pairs of coevolutionary profiles. This correlation value is added to the average of partial correlations of third proteins for each of the predictions to create an interaction score. The higher the interaction score, the more coevolutionary related these two proteins are and thus, the more likely it is that they are related functionally or structurally.

Finally, we observed that the predictions influenced by very few third proteins tended to be false positives (or true positives not recorded yet). To further increase the ranking power of this interaction score, a final update was introduced. A Mann-Whitney-U-Test was performed on the data and we found out that predictions influenced by a small number of third proteins are significantly more likely to be false positive (Statistic=23651311893.500, p=0.000). The rationale behind this penalization strategy is based on the understanding that highly specific protein-protein interactions (PPIs) should still reflect influence from other proteins within the same system. While the impact of third proteins on such PPIs may be weak, it should still be significant if the interaction occurs naturally. Conversely, PPIs that appear strongly correlated but show no significant influence from any third protein are unlikely to represent true biological interactions and are more likely artifacts of the method or other biases. This penalization approach adjusts for such cases by reducing the interaction score of protein pairs with no substantial third-protein influence (≤ 50 third proteins), effectively filtering out spurious predictions. The final list is then rearranged according to the Interaction Score (IS) considering this penalty, producing the final output of the ContextMirror updated implementation as follows:

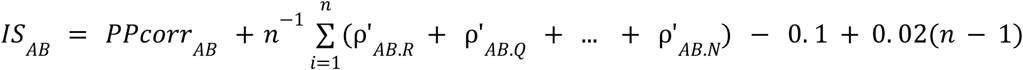

Where capital letters (*A*, *B*, *R*, *Q*, …, *N*) represent proteins, *n* is the number of significant third profiles influencing the interaction and *PPcorr_AB_* is the correlation value extracted from the profile-profile correlation matrix for the pair AB, and ρ’_*AB.N*_ is the partial correlation between the coevolutionary profiles of proteins A and B with, holding constant the coevolutionary profile of a third protein (e.g. N).

### 4.2 Analysis of the predicted PPIs

#### Benchmarking

As a benchmarking, we predicted the interactomes of three bacterial species from their reference proteomes: *Escherichia coli* (strain *K12 MG1655*), *Salmonella enterica* (subsp. enterica serovar *Typhimurium LT2*) and *Staphylococcus aureus* (strain *NCTC 8325* / *PS 47*), compared to the ground truth provided by experimental results stored in a combination of databases: IntAct ^53^, STRING ^54^, EcoCyc ^23^ and Interactome3D ^55^. Interactions with score below 0.2 were filtered out from the STRING database to improve the quality of the data. For *S. aureus*, due to insufficient strain-specific data, experimental information from experimental findings at the species level were also incorporated ^56^.

The precision is calculated as the number of true positive predictions divided by the number of total predictions (true positives and false positives). The recall is calculated as the number of true positive predictions divided by the number of relevant cases. For intra-community calculations, only edges contained in the retrieved communities are considered. For the combined network precision analysis in Figure 4 and **Table S2**, communities were automatically retrieved 100 times to calculate the mean and standard deviation. The selected communities in Figures 5**, 6, S5, S6, S7, S8, S9, S10** and **S11** were retrieved with the random seed that optimized the precision for shared predictions (seed = 76) as follows:

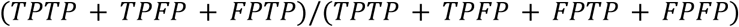

where the numerator represents any interaction that is a true positive in any of the datasets, and the denominator represents all predictions made in both datasets.

The chance precision for a given network is calculated as the number of true positive predictions divided by the number of potential pairs formed by all the nodes in the network (**Table S4**).

#### Dependency of the interaction score on MSA diversity

To assess the dependency of the interaction scores on the number of sequences contained in the paired MSAs, we quantified the sequence diversity of the MSAs defined by number of effective sequences (N_eff_) as follows:

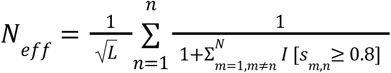

where *L* is the length of the query sequence, *N* is the number of sequences contained in the MSA, *S_m,n_* is the sequence identity between the *m*^th^ and *n*^th^ sequences, and *I*[.] represents the Iverson bracket, which takes the value *I*[*S_m,n_* ≥ 0.8] = 1 if *S_m,n_* ≥ 0.8, and 0 otherwise. After calculating N_eff_ for MSA A and MSA B in a given paired MSA, we defined the N_eff_ of the paired MSA as the minimum between N_eff_ A and N_eff_ B.

#### Structural analysis

For a number of predicted PPIs, structural models were generated using AlphaFold2.3.0 ^11,12^. For our analysis, we used the top-ranked unrelaxed model provided by AlphaFold. We adhered to the default settings for multiple sequence alignment (MSA) generation and the prescribed number of recycles. Our final dataset included 60 protein dimer complexes, with 20 each derived from *Escherichia coli*, *Staphylococcus aureus*, and *Salmonella enterica*. The selection criteria for the 20 proteins from each species was based on the interaction score. For each species, the top10 and bottom10 PPIs were selected. For each protein complex, we calculated the pDockQ2 score as detailed in ^12^, the improved version of pDockQ metric, designed to evaluate the quality of modelled protein complex models.

To evaluate the accuracy of the complex models, we identified the minimum pDockQ2 score for each protein pair (**Figure S12**). We adopted the threshold of 0.23, as recommended by the creators of pDockQ2, to signify a high-accuracy complex. Those complexes exceeding this threshold were chosen for detailed structural representation in Figure 2.

Full details regarding the protein complex IDs, their interaction scores, and the corresponding pDockQ2 scores are provided in **Table S5**.

#### Multispecies PPI networks

Based on the pairwise interaction scores outputted by CM2.0 we build a network in which each node represents an individual protein, and the edges denote the predicted interactions between them and are weighted by the interaction score. We conducted community searches using the Louvain algorithm recursively (resolution = 1) to identify protein communities (maximum community size = 30 nodes) either in individual or in combined networks. We also used the network to identify the sets of interactions made between certain proteins of interest in different species.

To enable comparisons between proteins from distinct proteomes, we identified the closest homologs in the other species. Using these homolog lists, proteins across species could be compared as considered equivalent when they are found to be a reciprocal best hit within the top 5 closest homologs for each pairwise comparison. For each of the proteins, a custom proteome database was created using the “makeblastdb” tool from the BLAST+ suite ^57^ and standard blastp parameters were used to retrieve homologs from each corresponding combination of species. **Table S6** gathers the results of these homolog lists.

For the combined networks of *E. coli* and *S. enterica*, nodes were superimposed according to the reciprocal best hit relationships calculated earlier. The color of the nodes indicate the network of origin, blue indicating *E. coli* and orange denoting *S. enterica*. Gray nodes represent reciprocal best hits. Edges follow the same logic and additionally, the edge can either be represented as a solid line (validated interaction) or a dashed line (unvalidated interaction). When predicted in both, the weight of the nodes is the average of the interaction scores predicted in both individual networks.

### 4.3 Technical specifications

#### Software requirements

We used BLAST version 2.13 to search the NCBI database for potential orthologs and MAFFT (Multiple Alignment using Fast Fourier Transform) version 7.471 to build MSAs ^58^. The specifications of the alignment design include the *automatic* parameter that automatically chooses the best strategy between progressive (FFT-NS-2) and iterative (FFT-NS-I or L-INS-i) methods, and the *anysymbol* option that considers ambiguous residues (e.g., B, Z, J, X, period, …). The Python version employed to run this program is 3.6.1 with the following dependencies: os, sys, Bio v1.79 (SeqIo, AlignIo, Phylo, ClustalWCommandline and DistranceTreeConstructor), numpy v1.21.5, networkx v2.8.5, requests v2.28.1, pandas v1.4.3, community v0.16 and scipy v1.8.1. Additionally, FastTree v2.1 and csvkit v1.3.0 are needed. Any parallelization tool (e.g. GREASY) is recommended to increase speed.

Many of these are novel additions to the pipeline, as numerous beneficial tools have been developed since the original method was published. These adaptations include the update of the alignment software from MUSCLE to MAFFT, an overall faster and more consistent method ^59^. Likewise, we replaced ClustalW with FastTree ^60^ as the phylogenetic-tree-building algorithm. Also, since the size of databases has increased drastically over the past years, the GREASY parallelization strategy was implemented to reduce the computing time and manage large datasets. Moreover, the update of the BLAST software and the local installation of databases significantly increased the speed of RBH operations.

Details regarding the resource usage and computing time allocated for predicting the three interactomes are presented in **Table S7**.

## Notes

### Competing Interest Statement

The authors have declared no competing interest.

