## Supplementary Information for "Sequence-based coevolutionary prediction of species-specific interactomes"

This PDF file includes:

- Figures S1 to S12
- Tables S1 to S7

### Supplementary Figures

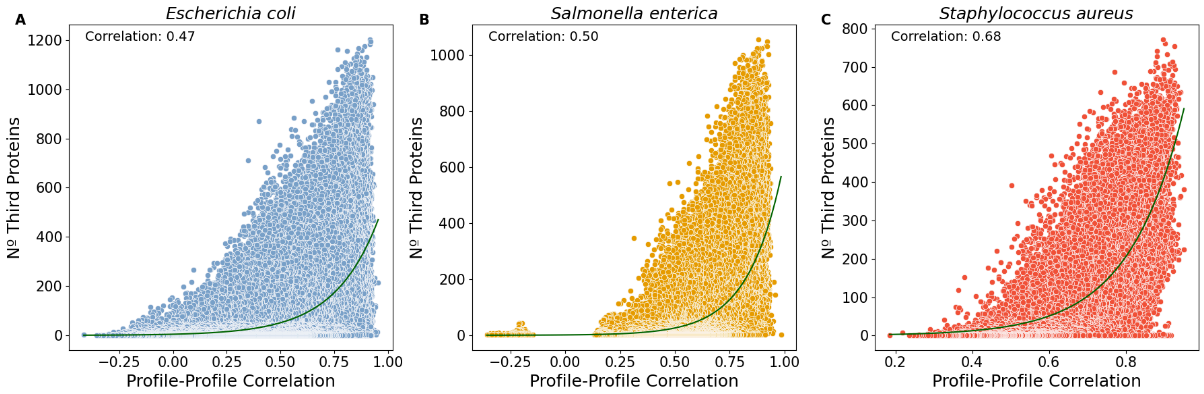


**Figure S1.** Correlation between the Profile-Profile Correlation and the number of third proteins for every predicted pair in *Escherichia coli* (A), *Salmonella enterica* (B) and *Staphylococcus aureus* (C). The green line represents the exponential curve fitted to the distribution of points in the scatter plot. The correlation displayed in the plots reflects the Pearson correlation between the Profile-Profile Correlation and the number of third proteins influencing each pair.

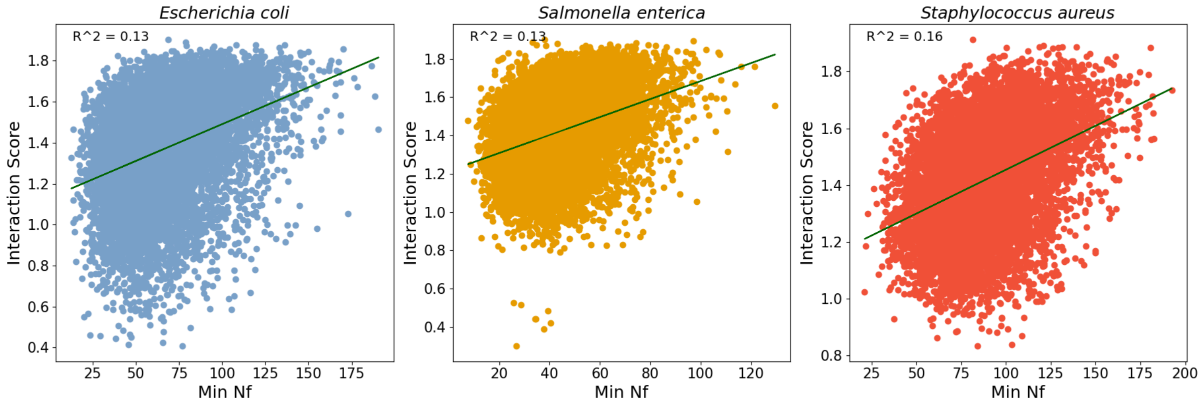


**Figure S2.** Correlation between the Interaction Score and the effective number of sequences (Neff) for 10000 random predicted pairs in *Escherichia coli* (A), *Salmonella enterica* (B) and *Staphylococcus aureus* (C). The green line represents the linear regression line fitted to the distribution of points in the scatter plot. The coefficient of determination ($R^2$) for each model is displayed for each case.

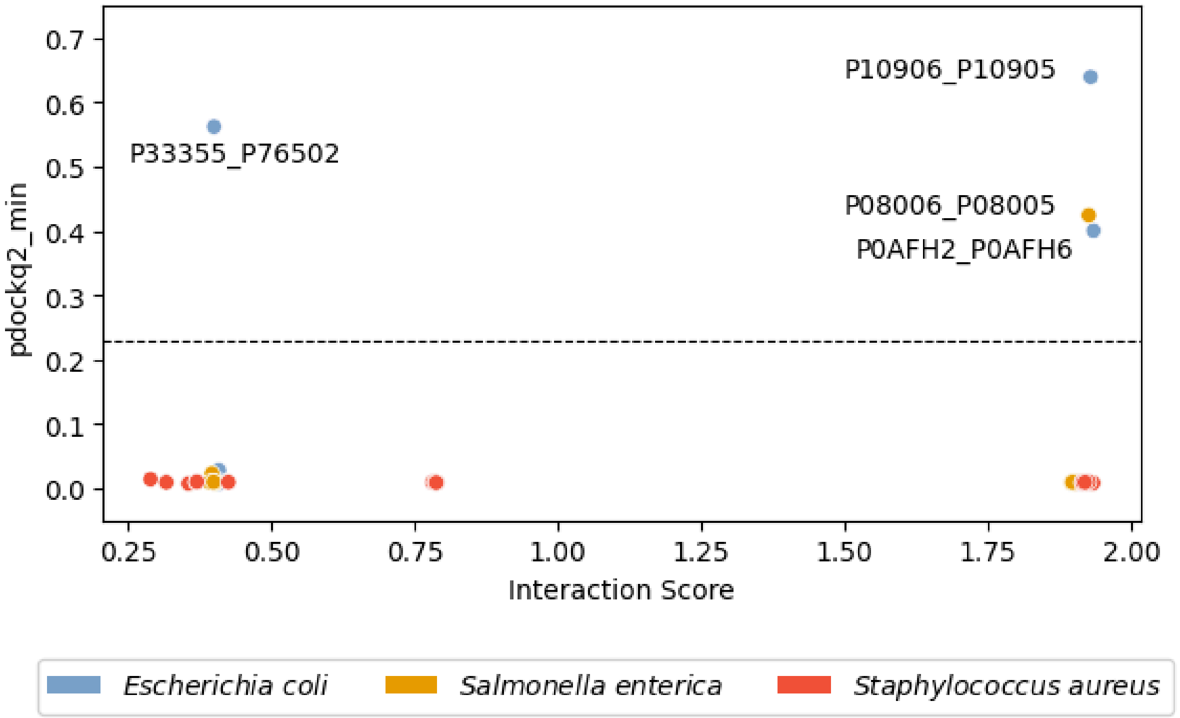


**Figure S3.** Results of minimum pDockQ2 score per PPI versus interaction score. Dashed line is the threshold of the pDockQ2 score (0.23) to be considered as a prediction of likely interaction. Predictions of AF Multimer for 20 proteins of *E. coli*, *S. aureus* and *S. enterica* each, 10 with high and 10 with low ContextMirror Interaction Score. Dataset S5 contains the data used to produce this plot.

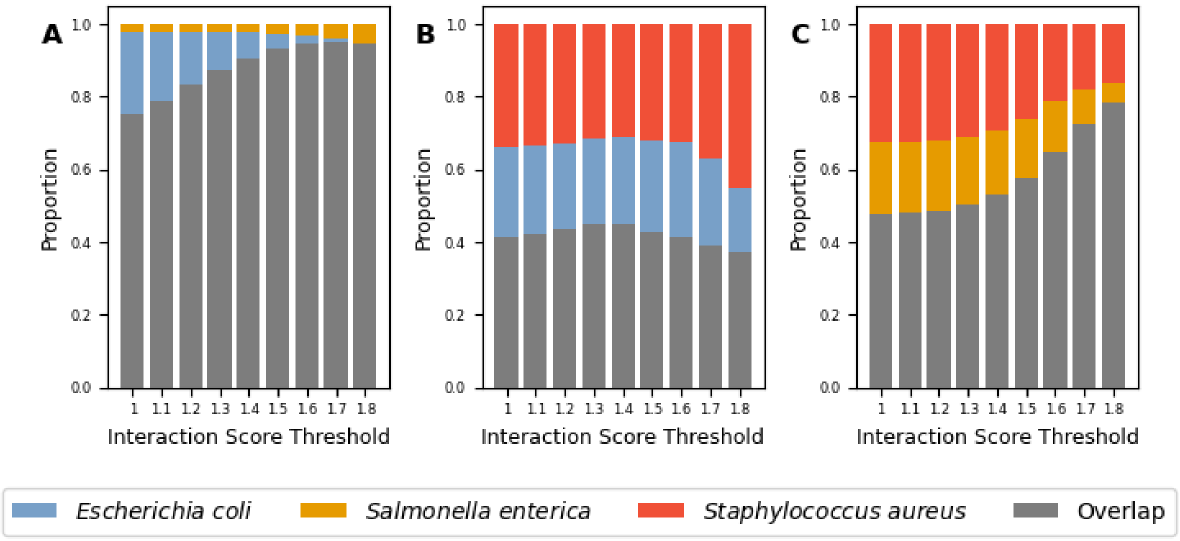


**Figure S4.** Proportion of shared PPIs at different interaction score thresholds for the three pairwise comparisons.

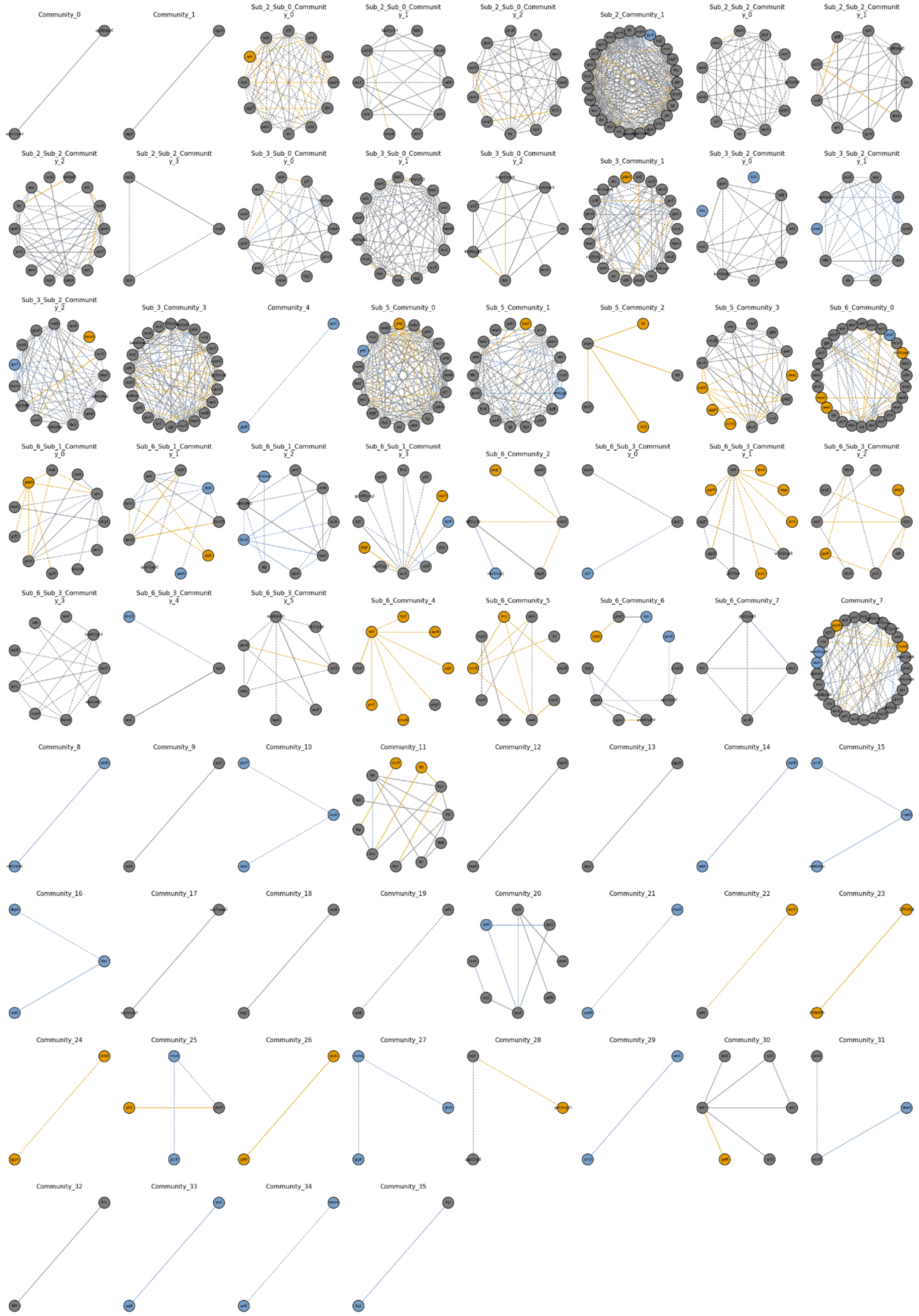


**Figure S5.** Communities retrieved using the Louvain algorithm on the union of the predicted interactomes for *E. coli* and *S. enterica* with an interaction score threshold of 1.8. Common nodes and edges are colored gray, blue indicates predictions exclusive of *E. coli* and orange shows unique predictions for *S. enterica*.

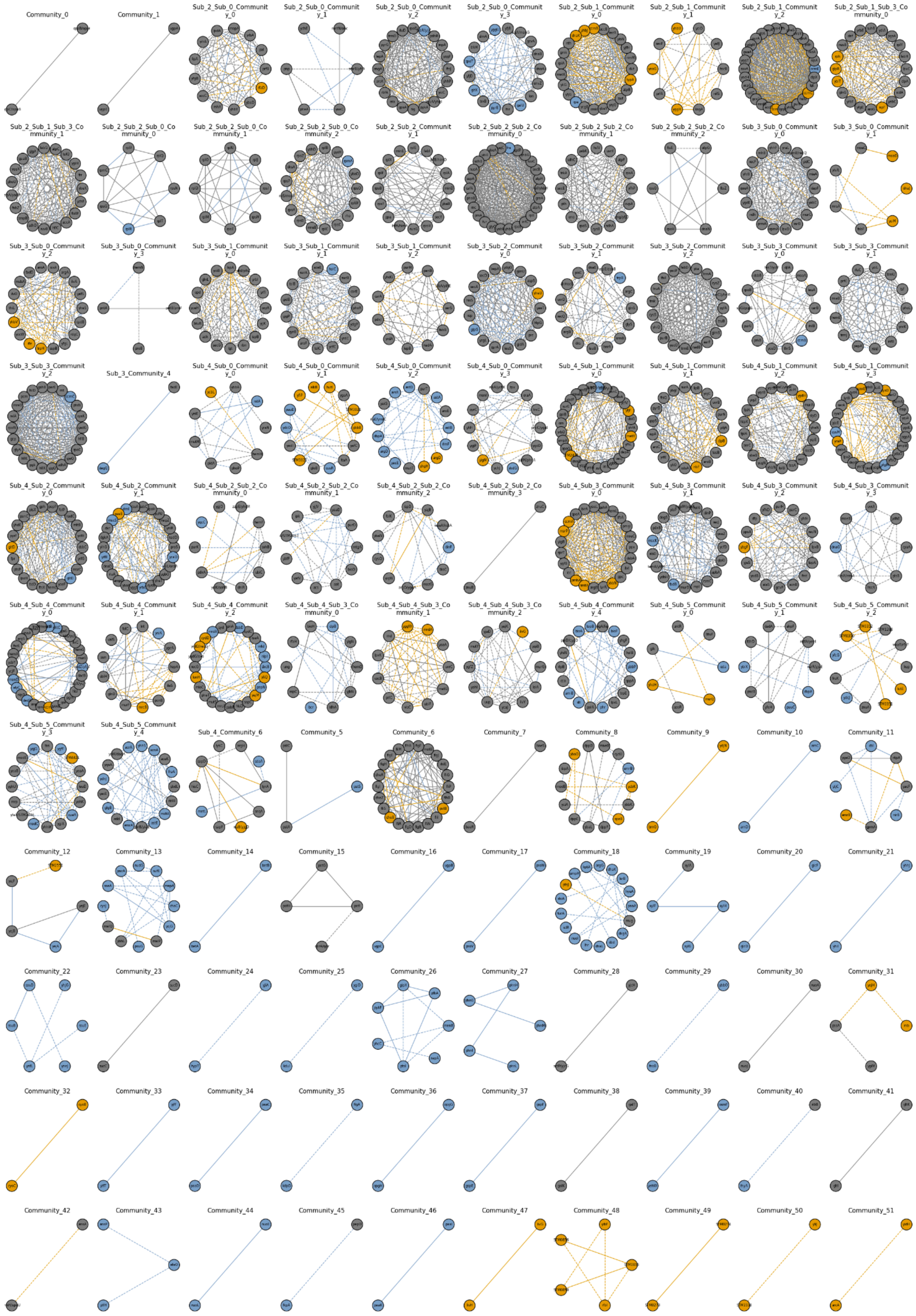

**Figure S6.** Communities retrieved using the Louvain algorithm on the union of the predicted interactomes for *E. coli* and *S. enterica* with an interaction score threshold of 1.7. Common nodes and edges are colored gray, blue indicates predictions exclusive of *E. coli* and orange shows unique predictions for *S. enterica*.

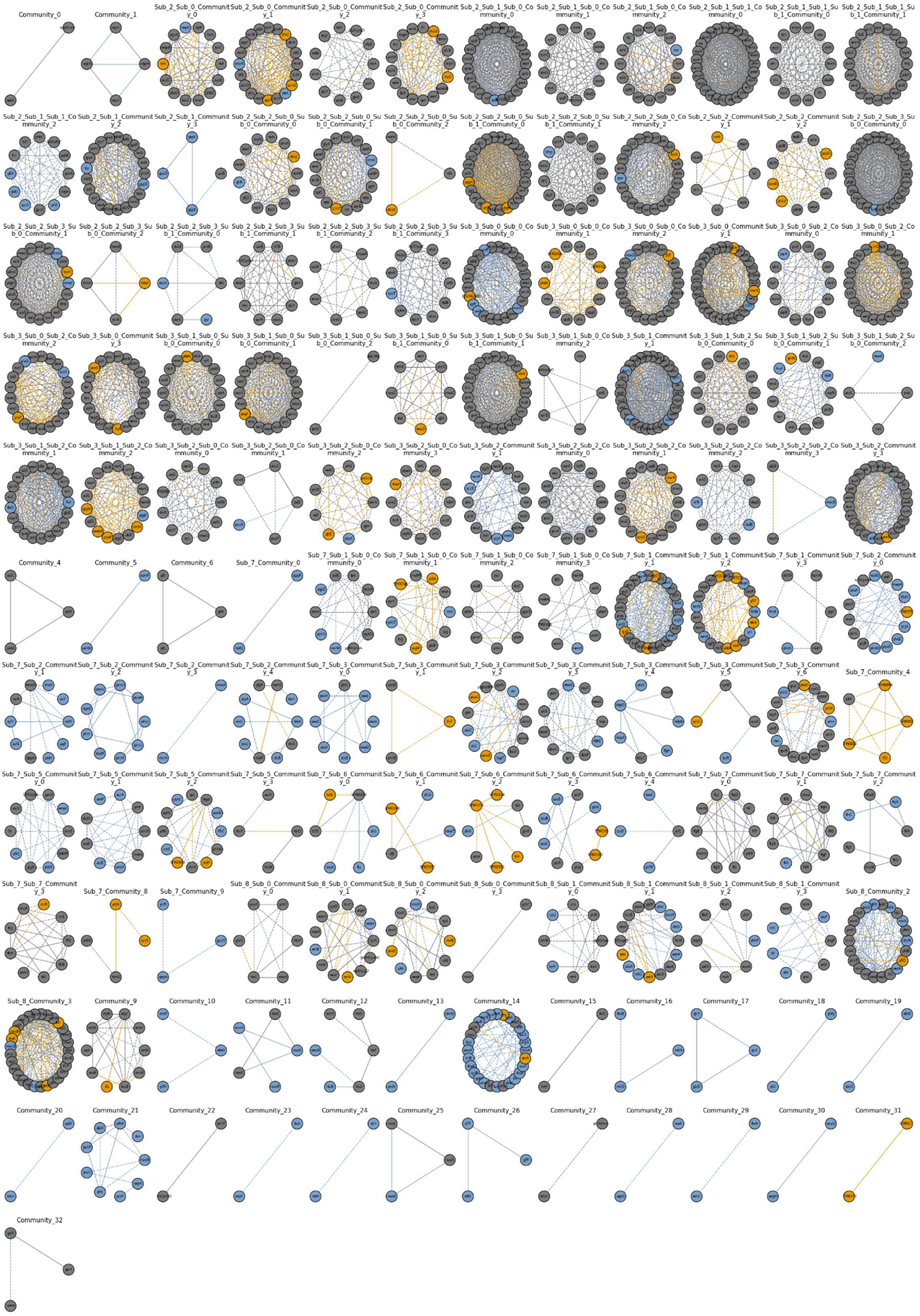

**Figure S7.** Communities retrieved using the Louvain algorithm on the union of the predicted interactomes for *E. coli* and *S. enterica* with an interaction score threshold of 1.6. Common nodes and edges are colored gray, blue indicates predictions exclusive of *E. coli* and orange shows unique predictions for *S. enterica*.

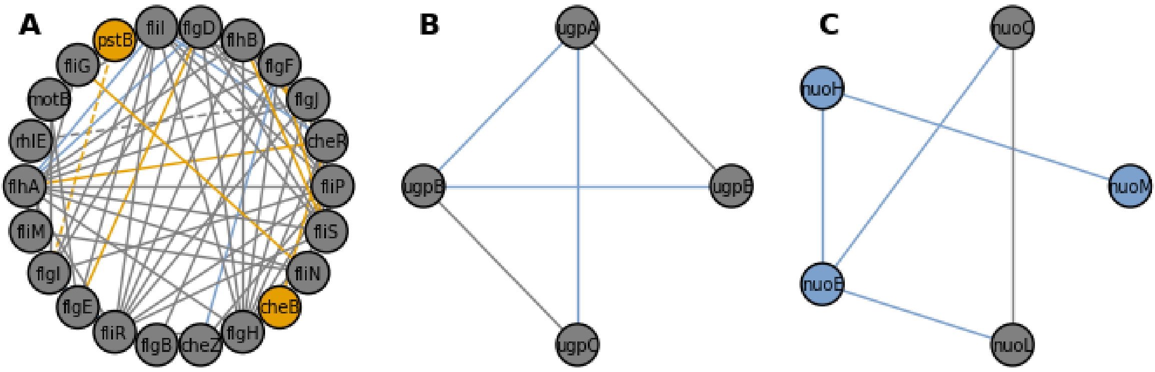


**Figure S8.** Functional communities retrieved from the combined network of *E. coli* and *S. enterica* predicted interactomes at different interaction score thresholds. A) Flagellar assembly complex (IS > 1.7). B) ABC transporter complex UgpBAEC (IS > 1.6). C) NADH:ubiquinone oxidoreductase complex (IS > 1.6).

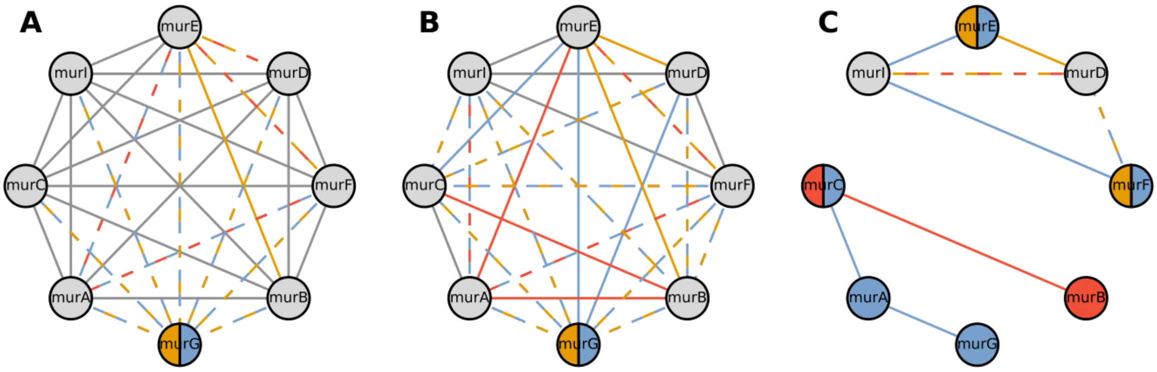


**Figure S9**. Predicted protein-protein interactions (PPIs) for the Mur complex in *E. coli, S. enterica,* and *S. aureus*, filtered with interaction thresholds of 1.2 (**A**), 1.5 (**B**), and 1.7 (**C**). Shared nodes and edges are depicted in gray, *E. coli* predictions are shown in blue, *S. enterica* predictions are highlighted in orange, and *S. aureus* predicted interactions are indicated in red.

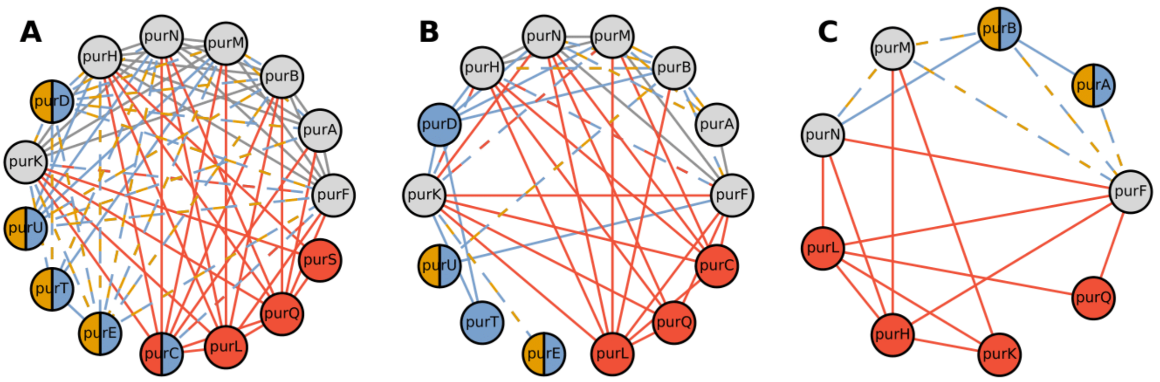


**Figure S10**. Predicted protein-protein interactions (PPIs) for the Pur complex in *E. coli, S. enterica*, and *S. aureus*, filtered with interaction thresholds of 1.2 (**A**), 1.5 (**B)**, and 1.7 (**C**). Shared nodes and edges are depicted in gray, *E. coli* predictions are shown in blue, *S. enterica* predictions are highlighted in orange, and *S. aureus* predicted interactions are indicated in red.

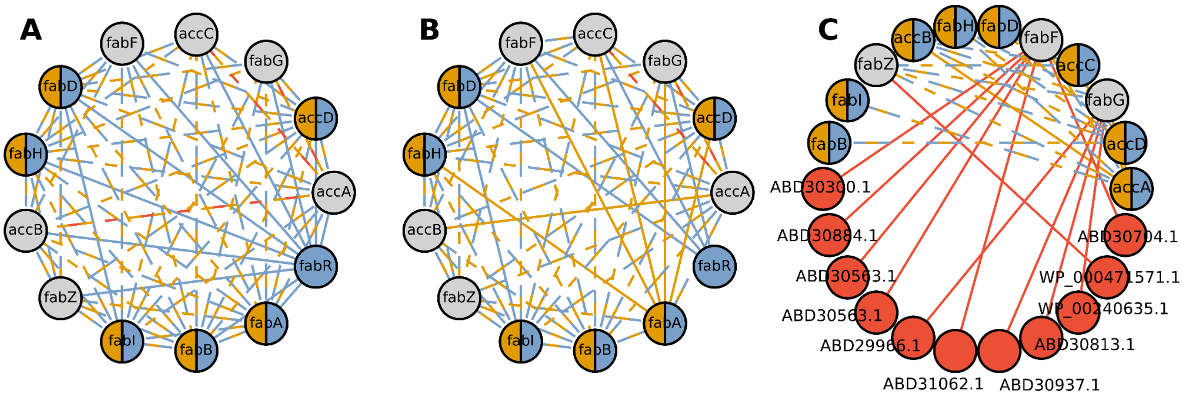


**Figure S11**. Predicted protein-protein interactions (PPIs) for the acc+fab complex in *E. coli, S. enterica*, and *S. aureus*, filtered with interaction thresholds of 1.5 (**A**), 1.6 (**B**). (**C**) Predicted protein-protein interactions (PPIs) for the acc+fab complex in *E. coli, S. enterica*, and *S. aureus*, filtered with interaction thresholds of 1.7, including unmapped proteins in *S. aureus* . Shared nodes and edges are
depicted in gray, *E. coli* predictions are shown in blue, *S. enterica* predictions are highlighted in orange, and *S. aureus* predicted interactions are indicated in red.

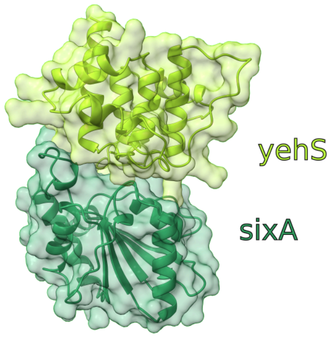


**Figure S12.** AlphaFold Multimer model of P33355 - P76502 (yehS - sixA) validated structurally (min pDockQ2 = 0.562067), but predicted to have a low interaction score (IS = 0.40), yet not reported as an interaction in *E. coli* (TN).

### Supplementary Tables

**Table S1.** Pairwise precision at different interaction score (IS) thresholds for different interactome predictions at various interaction score thresholds.

| | IS > 1.8 | IS > 1.7 | IS > 1.6 |
|---|---|---|---|
| *E. coli* | 34,8% | 25,7% | 20,8% |
| *S. enterica* | 34,6% | 25,7% | 20,6% |
| *S. aureus* | 41.40% | 30.14% | 25.41% |

**Table S2.** Summary of predictions included in communities extracted automatically from the union of the predicted interactomes of *E. coli* and *S. enterica* with three interaction score thresholds (Figures S5-S7). GT stands for ground truth and IS is the interaction score from ContextMirror2.0.

| **Predicted in** | **GT *E. coli*** | **GT *S. enterica*** | **PPIs (IS > 1.6)** | **Std. Dev. (IS > 1.6)** | **PPIs (IS > 1.7)** | **Std. Dev. (IS > 1,7)** | **PPIs (IS > 1.8)** | **Std. Dev. (IS > 1.8)** |
|---|---|---|---|---|---|---|---|---|
| **Both** | **TP** | **TP** | 900,56 | 40,79 | 641,02 | 30,25 | 307,94 | 28,38 |
| **Both** | **TP** | **FP** | 326,35 | 19,62 | 213,31 | 15,91 | 104,11 | 10,6 |
| **Both** | **FP** | **TP** | 293,59 | 18,24 | 186,30 | 10,18 | 77,74 | 8,29 |
| **Both** | **FP** | **FP** | 3.398,19 | 167,32 | 1.892,86 | 87,95 | 610,10 | 43,82 |
| ***E. coli*** | **TP** | **-** | 263,86 | 11,9 | 174,43 | 11,34 | 86,16 | 7,14 |
| ***E. coli*** | **FP** | **-** | 1.088,67 | 43,68 | 592,11 | 24,76 | 163,03 | 11,75 |
| ***S. enterica*** | **-** | **TP** | 136,34 | 9,03 | 102,03 | 9,10 | 61,07 | 6,5 |
| ***S. enterica*** | **-** | **FP** | 781,33 | 48,87 | 482,10 | 29,76 | 164,93 | 14,28 |
| Precision *E. coli* | | | **23,77** | **-** | **27,8** | **-** | **36,93** | **-** |
| Precision *S. enterica* | | | **22,8** | **-** | **26,42** | **-** | **33,7** | **-** |

**Table S3.** PPI comparative metrics across reference proteomes of *E. coli, S. enterica,* and *S. aureus.*

| Organism | *Escherichia coli* | *Salmonella enterica* | Staphylococcus aureus |
|---|---|---|---|
| Nº proteins in reference proteome | 4298 | 4814 | 2889 |
| Nº combinations (PPIs) | 8 944 334 | 10 771 760 | 3 689 685 |
| Nº pairs (>1000 common species) | 718073 | 634530 | 327697 |
| Nº pairs significantly correlated | 542348 | 377123 | 121738 |
| Ground truth (nº PPI) | 584 411 | 491 135 | 107 559 |

**Table S4.** Calculated chance precision metrics for predicted PPIs in *E. coli, S. enterica,* and *S. aureus*

| Organism | Nº True Positives | Nº proteins (N) | Potential PPIs N*(N-1)/2 | Random Chance (Fraction) | Random Chance (Percentage) |
|---|---|---|---|---|---|
| *E. coli* | 312,961 | 4126 | 8,509,875 | 312,961 / 8,509,875 | 3.6% |
| *S. enterica* | 295,678 | 4447 | 9,885,681 | 295,678 / 9,885,681 | 2.9% |
| *S. aureus* | 67,737 | 1481 | 3,065,670 | 67,737 / 3,065,670 | 2.2% |

**Table S5**. Summary of protein complexes predicted using AlphaFold Multimer v2.3, with CM2.0 scores and pDockQ2 quality metrics. The table presents organisms, their associated complexes, CM2.0 scores, true positive (TP) validation status, and the minimum pDockQ2 scores for each complex. The pDockQ2 metric evaluates the quality of individual interfaces within a multimer, with the lowest pDockQ2 score for each complex shown. The TP column indicates either "NO" or the name of the source where validation was obtained. Rows in bold highlight complexes with a minimum pDockQ2 score exceeding 0.23, indicating higher interface quality.

| Organism | Complex | CM2.0 score | TP | Min. pDockQ2 |
|---|---|---|---|---|
| **E. coli** | **P0AFH2_P0AFH6** | **1.934** | **ECOCYC_COMPLEXES** | **0.401** |
| **E. coli** | **P10906_P10905** | **1.929** | **ECOCYC_COMPLEXES** | **0.639** |
| E. coli | P0A6P5_P0A8I1 | 1.906 | STRING | 0.008 |
| E. coli | P0AG86_P0ABA6 | 1.904 | NO | 0.01 |
| E. coli | P33643_P0A6P7 | 1.904 | STRING | 0.009 |
| E. coli | P06616_P0A9J0 | 1.904 | NO | 0.009 |
| E. coli | P33643_P23839 | 1.901 | NO | 0.009 |
| E. coli | P0AFF6_P0ABA4 | 1.901 | STRING | 0.009 |
| E. coli | P21599_P33643 | 1.9 | NO | 0.009 |
| E. coli | P0AFF6_P0A9J0 | 1.9 | NO | 0.009 |
| **E. coli** | P0A850_P0A8T5 | 0.409 | NO | 0.028 |
| E. coli | P0AB18_P69913 | 0.409 | NO | 0.012 |
| **E. coli** | P0ABF8_P69913 | 0.408 | NO | 0.007 |
| E. coli | P0A9B2_P0A9H9 | 0.407 | STRING | 0.009 |
| E. coli | P25745_P33355 | 0.405 | NO | 0.009 |
| E. coli | P21645_P69913 | 0.403 | STRING | 0.009 |
| E. coli | P33355_P04425 | 0.402 | NO | 0.01 |
| **E. coli** | **P33355_P76502** | **0.4** | **NO** | **0.562** |
| E. coli | P0A7F9_P69913 | 0.4 | NO | 0.009 |
| E. coli | P0A9K3_P0A6V1 | 0.398 | NO | 0.009 |
| S. aureus | mfd_rqcH | 1.934 | NO | 0.008 |

| | | | | |
|---|---|---|---|---|
| S. aureus | mfd_alaS | 1.93 | YES | 0.008 |
| S. aureus | gpsA_plsX | 1.921 | YES | 0.007 |
| S. aureus | yycJ_mnmE | 1.92 | NO | 0.009 |
| S. aureus | mutL_priA | 1.917 | YES | 0.01 |
| S. aureus | prkC_polA | 1.916 | NO | 0.009 |
| S. aureus | rqcH_pcrA | 1.915 | NO | 0.009 |
| S. aureus | priA_polA | 1.914 | YES | 0.009 |
| S. aureus | hprK_WP_000838485 | 0.789 | unmapped | 0.009 |
| S. aureus | rplA_WP_000838485 | 0.788 | unmapped | 0.009 |
| S. aureus | nfo_WP_000838485 | 0.786 | unmapped | 0.009 |
| S. aureus | yurK_ABD29612 | 0.425 | unmapped | 0.01 |
| S. aureus | yurK_yabD | 0.37 | NO | 0.01 |
| S. aureus | yurK_gyrB | 0.37 | NO | 0.009 |
| S. aureus | yurK_atpD | 0.356 | NO | 0.007 |
| S. aureus | yurK_yvcJ | 0.317 | NO | 0.009 |
| S. aureus | gatA_WP_000838485 | 0.781 | unmapped | 0.009 |
| S. aureus | mnmG_WP_000229248 | 1.92 | unmapped | 0.009 |
| S. aureus | tig_WP_000229248 | 1.925 | unmapped | 0.01 |
| S. aureus | yurK_rmnV | 0.29 | NO | 0.014 |
| **S. enterica** | **P08006_P08005** | **1.926** | **YES** | **0.424** |
| S. enterica | Q7CQX1_Q7CPM4 | 1.904 | YES | 0.01 |
| S. enterica | Q7CPZ7_Q7CPD5 | 1.902 | YES | 0.009 |
| S. enterica | Q56057_Q7CPM4 | 1.902 | YES | 0.008 |
| S. enterica | Q56016_Q7CQE8 | 1.9 | YES | 0.009 |
| S. enterica | Q7CPM4_Q8ZL95 | 1.899 | NO | 0.01 |
| S. enterica | Q7CPJ7_Q8ZKW8 | 1.899 | YES | 0.009 |
| S. enterica | Q56016_P0A287 | 1.899 | YES | 0.009 |
| S. enterica | Q8ZNB9_P40731 | 1.898 | YES | 0.009 |

| | | | | |
|---|---|---|---|---|
| S. enterica | Q7CQX1_Q8ZL95 | 1.896 | NO | 0.009 |
| S. enterica | P07801_Q7CPZ5 | 0.404 | NO | 0.01 |
| S. enterica | P07801_P65774 | 0.402 | NO | 0.009 |
| S. enterica | P69917_Q8ZMB8 | 0.401 | NO | 0.009 |
| S. enterica | Q8ZM48_Q7CPQ2 | 0.401 | NO | 0.01 |
| S. enterica | Q8ZN19_P69917 | 0.399 | NO | 0.01 |
| S. enterica | P69917_Q8ZLZ4 | 0.399 | YES | 0.009 |
| S. enterica | P67073_P69917 | 0.398 | NO | 0.009 |
| S. enterica | P63866_P60446 | 0.396 | NO | 0.023 |
| S. enterica | O85140_P69917 | 0.396 | NO | 0.01 |
| S. enterica | Q8ZM48_Q8ZLS7 | 0.393 | NO | 0.009 |

**Table S6.** Reciprocal best hits (RBH) considering the top 5 homolog results for each potential comparison. n indicates the size of each proteome.

| | ***E. coli*** | ***S. enterica*** | ***S. aureus*** |
|---|---|---|---|
| *E. coli* (n = 4298) | - | 3583 | 2069 |
| *S. enterica* (n = 4814) | - | - | 2245 |
| *S. aureus* (n = 2786) | - | - | - |

**Table S7.** Computational time and resource allocation for ContextMirror2.0.

| | ***Escherichia coli*** | ***Salmonella enterica*** | ***Staphylococcus aureus*** | **Average** |
|---|---|---|---|---|
| Estimated total time (h) on a single CPU | 110 000 | 86 845.5 | 65 041.5 | 87 296 ± 22 482 |
| Estimated time / PPI (s) on a single CPU | 44 | 29 | 63 | 45 ± 17 |
| Time to process 10 000 PPIs using 100 CPUs | 1h 14m | 49m | 1h 46m | 1h 16m ± 28m |
